# Metabolic Modeling Elucidates the Transactions in the Rumen Microbiome and the Shifts upon Virome Interactions

**DOI:** 10.1101/703843

**Authors:** Mohammad Mazharul Islam, Samodha C. Fernando, Rajib Saha

**Affiliations:** Department of Chemical and Biomolecular Engineering, University of Nebraska-Lincoln, Lincoln, NE-68588, USA; Department of Animal Science, University of Nebraska-Lincoln, Lincoln, NE-68588, USA

**Keywords:** microbial community, microbiome-virome interaction, rumen, genome-scale metabolic modeling, viral auxiliary metabolic genes

## Abstract

The complex microbial ecosystem within the bovine rumen plays a crucial role in host nutrition, health, and environmental impact. However, little is known about the interactions between the functional entities within the system, which dictates the community structure and functional dynamics and host physiology. With the advancements in high-throughput sequencing and mathematical modeling, *in silico* genome-scale metabolic analysis promises to expand our understanding of the metabolic interplay in the community. In an attempt to understand the interactions between microbial species and the phages inside rumen, a genome-scale metabolic modeling approach was utilized by using key members in the rumen microbiome (a bacteroidete, a firmicute, and an archaeon) and the viral phages associated with them. Individual microbial host models were integrated into a community model using multi-level mathematical frameworks. An elaborate and heuristics-based computational procedure was employed to predict previously-unknown interactions involving the transfer of fatty acids, vitamins, coenzymes, amino acids, and sugars among the community members. While some of these interactions could be inferred by the available multi-omic datasets, our proposed method provides a systemic understanding of why these occur and how these affect the dynamics in a complex microbial ecosystem. To elucidate the functional role of the virome on the microbiome, local alignment search was used to identify the metabolic functions of the viruses associated with the hosts. The incorporation these functions demonstrated the role of viral auxiliary metabolic genes in relaxing the metabolic bottlenecks in the microbial hosts and complementing the inter-species interactions. Finally, a comparative statistical analysis of different biologically significant community fitness criteria identified the variation in flux space and robustness of metabolic capacities of the community members. Our elucidation of metabolite exchange among three members of rumen microbiome shows how their genomic differences and interactions with the viral strains shape up a highly sophisticated metabolic interplay and explains how such interactions across kingdoms can cause metabolic and compositional shifts in the community and affect the health, nutrition, and pathophysiology of the ruminant animal.

## Introduction

Within the ruminant species, the microbial community has co-evolved with its host and has helped the host animal obtain energy from low quality fiber rich diets (Hungate, 1975;Hobson, 1988;Henderson et al., 2015). The feed ingested by the ruminant animal undergoes extensive microbial digestion and fermentation in the cattle rumen, producing a range of short chain fatty acids (SCFAs) and energy for the host, and also releases methane, hydrogen, and carbon di-oxide to the atmosphere (Nocek and Russell, 1988). This anaerobic environment is densely (>10^10^–10^11^ cells/g of contents) populated by diverse and interdependent species of Bacteria, Protozoa, Fungi, Archaea and Viruses, which are involved in breakdown of complex carbohydrates and polymers from plants, hydrogen transfer, and inter-species transaction of fermentation products and of oligomers and monomers (Bryant and Burkey, 1953;Hungate, 1966;Russell and Rychlik, 2001;Flint et al., 2008). Among the major bacterial players are the phyla Bacteroidetes and Firmicutes. Majority of the Bacteroidetes (specifically *P. ruminicola* and *P. bryantii*) have a significant role in breakdown of starch and xylan polysaccharides and in the metabolism of proteins and peptides (Wallace et al., 1997;Flint et al., 2008;Thomas et al., 2011). Additionally, Firmicutes including *Butyrivibrio fibrisolvens, Ruminococcus flavefaciens, Ruminococcus albas, Eubacterium cellulosolvens* etc. play an essential role in the metabolism of cellulose (Flint et al., 2008;Henderson et al., 2015). Although not highly abundant, methanogenic Archaea are also present within the rumen and many of the methanogens (including *Methanobrevibacter gottschalkii, Methanobrevibacter ruminantium, Methanosarcina barkeri, Methanosarcina mazeii* etc.) are involved in methanogenesis (St-Pierre and Wright, 2013;Henderson et al., 2015;Seedorf et al., 2015;Danielsson et al., 2017). Methanogens are responsible for the release of methane and other gases in the atmosphere while consuming SCFAs, carbon di-oxide, and hydrogen from other microbes including Bacteroidetes or Firmicutes. The stability and diversity of the rumen microbiome is critical for the animals’ health, nutrition, immunity, and survival. Prior studies concluded that disruption in the community composition by sudden administration of grain or glucose to a ruminant animal previously on a dried forage ration often leads to the damage of the rumen tissues and death of the animal within as short of time as 18 hr due an explosive growth of the Firmicute *Streptococcus bovis* and *Lactobacillus spp*., and accumulation of abundant lactic acid (Hungate et al., 1952;Owens et al., 1998;Russell and Rychlik, 2001), thus causing lactic acidosis. In addition, methane production in cattle correlates with rumen methanogenic Archaea and bacterial community structure and dietary composition (Moss et al., 2000;Danielsson et al., 2017). Hence, rumen microbial community remains one of the most interesting and poorly explored natural ecosystems.

The complex interactions between bacterial host and viral phages associated with them drive the ecological dynamics and behavior in many natural systems (Lenski, 1988). Viral modifications of microbial and cyanobacterial (Thompson et al., 2011;Hurwitz and U’Ren, 2016) metabolism was identified in a substantial number of natural systems including marine ecosystem, infectious human diseases, aquifer sediments, and animal gut ecosystems (Sullivan et al., 2006;Suttle, 2007;Minot et al., 2011;Hurwitz et al., 2013;Pan et al., 2014;Weitz et al., 2015;Crummett et al., 2016;De Smet et al., 2016;Howe et al., 2016). Viruses affect intestinal and ruminal microbial ecosystem in cattle through a myriad of processes including cell lysis, energy production, reproduction, and reprogramming of microbial metabolism via Auxiliary Metabolic Genes or AMGs (Berg Miller et al., 2012;Parmar et al., 2016). Studies in recent years have demonstrated that viral AMGs augment the breakdown of complex plant carbohydrates and boost energy production and harvest, while accelerating viral replication inside the host (Anderson et al., 2017). However, a complete understanding of the complex virome-microbiome interaction and the roles of AMGs in the metabolic reprogramming of the host is still in its infancy.

Although advancements in high-throughput sequencing provide access to the vast diversity and makeup of this complex microbial ecosystem, our understanding of the factors that shape rumen microbial communities and interactions among them is rudimentary at best, and little is known about the processes shaping the distribution of rumen viruses or the modulation of microbe-driven processes in the rumen. In addition, the study of the rumen ecosystem suffers due to the lack of truly selective media or unique end products, which renders rapid identification of the community composition and metabolic states nearly impossible (Hungate, 1966;Krause and Russell, 1996a;Krause and Russell, 1996b). The study of ruminal ecology is further complicated by the observation that approximately 75% of the ruminal bacteria are tightly attached to feed particles or are found in biofilms (Hungate, 1966) and cannot be analyzed by only studying fecal contents. On the other hand, many of the virome-microbiome interactions studies appear to be focused on interactions between infectious human viruses and bacteria in an effort to understand respiratory infectious diseases (Levin et al., 1977;Pettigrew et al., 2011;Bosch et al., 2013;Opatowski et al., 2013;Shrestha et al., 2013). A recent study of virus-bacterial interactions in the rumen identified approximately 28,000 viral sequences present in the rumen, which found that the majority of viruses belong to a diversity of viral families including *Siphoviridae, Myoviridae*, and *Podoviridae* interacting with host bacterial phyla such as Firmicutes and proteobacteria (Berg Miller et al., 2012).

While complex biological systems are often challenging to be clearly understood or deciphered, explicit mathematical relation-based computational modeling promise *in silico* evaluation and analysis of the biological phenomena. With the gradual increase in computational capacity and the abundance of *in silico* genome-scale metabolic reconstruction tools, metabolic network models combined with constraint-based analysis provide a host of methods to explore, make discovery and hypotheses, and redesign biological systems at a genome-level (Varma and Palsson, 1994;Mahadevan et al., 2002;Papin et al., 2002;Burgard and Maranas, 2003b;Price et al., 2003;Palsson, 2006;Feist et al., 2009;Oberhardt et al., 2009b;Feist and Palsson, 2010;Henry et al., 2010;Orth et al., 2010;Schellenberger et al., 2010;Thiele and Palsson, 2010;Zomorrodi et al., 2012;Maranas and Zomorrodi, 2016;Islam and Saha, 2018). Genome-scale metabolic modeling offers the opportunity not only to map the metabolic landscape of single organisms but also to explore microbe-microbe and phage-microbe interactions. A number of computational tools were developed to model the interactions and dynamics in multi-species microbial communities in the past years (Stolyar et al., 2007;Salimi et al., 2010;Bordbar et al., 2011;Tzamali et al., 2011;Zhuang et al., 2011;Zhuang et al., 2012;Zomorrodi and Maranas, 2012;Zomorrodi et al., 2014;Henry et al., 2016;Mendes-Soares et al., 2016;Chan et al., 2017;Hanemaaijer et al., 2017;Mendes-Soares and Chia, 2017;Zomorrodi and Segre, 2017;Zuniga et al., 2017;Zengler and Zaramela, 2018). For modeling of multi-species communities, extensive experimental data on different omics’ level and a significant knowledge of the inter-species interactions are needed. However, the knowledge-base for microbe-microbe as well as host-microbe interactions is still very poor, and cross-feeding and/or interaction experiments cannot yet be routinely carried out *in silico* (Fritz et al., 2013). Thus, a combined computational-experimental approach can accelerate new discoveries in the realm of microbe-microbe and microbe-host metabolic interactions. Despite the limitations of current *in silico* reconstructed host-microbe interaction models, such efforts are of utmost importance because they allow for a detailed metabolic resolution of the complex relationships within microbial communities and with their host. Recently, Heinken and co-workers reconstructed and analyzed the first integrated stoichiometric model of murine and *Bacteroides thetaiotaomicron* metabolism and demonstrated the beneficial interaction of the host and the commensal microbes in the gut (Heinken et al., 2013). There have been several efforts to reconstruct individual and integrated community models of human gut microbes in recent years, using representative species from dominant classes of microorganisms (Heinken et al., 2013;Shoaie et al., 2013). However, a genome-scale metabolic analysis of the complex community in the rumen was never attempted before.

We developed a simplified and representative rumen community metabolic model with *Ruminococcus flavefaciens, Prevotella ruminicola*, and *Methanobrevibacter gottschalkii* as representative organisms from the three major functional guilds in the rumen ecosystem (*i.e*., Firmicutes, Bacteroidetes, and Archaea, respectively) mentioned above. These three organisms are responsible for fiber digestion, starch and protein digestion, and methane production, respectively. We reconstructed the draft models for each of these species by using the ModelSEED database (Henry et al., 2010), and then performed extensive manual curation, including chemical and charge-balancing, eliminating thermodynamically infeasible cycles, and ensuring network connectivity. The curated models of and *R. flavefaciens* (467 genes, 1033 metabolites, 1015 reactions), *P. ruminicola* (546 genes, 1069 metabolites, 1088 reactions), and *M. gottschalkii* (319 genes, 900 metabolites, 847 reactions) were integrated into a community model using a multi-level optimization framework (Zomorrodi and Maranas, 2012;Zomorrodi et al., 2014). The community model was used to estimate metabolite secretion profiles and community compositions. To enrich our understanding of the inter-species interactions in the ecosystem, we employed a detailed and comprehensive heuristic procedure that utilized existing GapFind-GapFill tools (Satish Kumar et al., 2007) and a subsequent series of knowledgebase-driven validations. We identified 22 novel interactions involving the transfer of fatty acids, vitamins, coenzymes, amino acids, and sugars among the community members. In addition, we bridged the network gaps in the pentose phosphate pathway, amino acid synthesis and utilization, nucleotide synthesis and degradation, purine metabolism, glycerophospholipid metabolism, and starch metabolism in the metabolic models of these organisms. To elucidate the functional role of the virome on the microbial ecosystem, we used local alignment search and identified metabolic functions of the viruses associated with the community members that drive nucleotide synthesis, reducing power generation, the reprogramming of the bacterial carbon metabolism to pentose phosphate pathway and folate biosynthesis, and viral replication. The identified functions of viral AMGs were incorporated into the model as additional metabolic functions. The addition of viral functionalities resulted in significant changes in bacterial metabolism, including relaxing metabolic bottleneck in the models, complementing microbe-microbe interactions, utilizing nutrients more efficiently and energy harvest by the host. We validated our results based on meta-transcriptomics, meta-proteomics and metabolomics studies on the rumen published recently (Saleem et al., 2012;Saleem et al., 2013;Li and Guan, 2017;Wang et al., 2019),

Overall, these findings support the hypothesis that viral AMGs play a crucial role in enhancing host fitness and robustness. We also studied the effect of using different community-level objective functions (*i.e*., growth, short-chain fatty acids production, plant feed utilization, greenhouse gas release, and small sugar molecule production) on the metabolic capacity of the community members. We found that the flux ranges of the microbial species are robust irrespective of the choice of a community objective. Hence, maximizing the community biomass is a rational choice since a stable community in the rumen needs to survive and grow at a reasonable rate to perform its necessary role in host nutrition and pathophysiology despite constant washout events like fecal secretion.

## Results

### Genome-scale models of *R. flavefaciens, P. ruminicola*, and *M. gottschalkii*

Three genome-scale metabolic models of representative organisms of the rumen microbiome were reconstructed for *P. ruminicola, R. flavefaciens*, and *M. gottschalkii* using the Modelseed database (Henry et al., 2010). This was followed by manual editing, refinements and further curation of the models. The draft reconstructions contained network gaps and 3-25% of reactions with chemical imbalances. One of them (*R. flavefaciens*) had thermodynamically inconsistent production of redox cofactors, which warranted an extensive manual curation to be performed on the models (see methods section for details). The manual curation steps ensured that there is no chemical or charge imbalance present in the models. The numbers of reactions that carries unrealistically high fluxes without any nutrient uptake (thus defined as reactions with unbounded fluxes) also decreased substantially (79% for *R. flavefaciens*, 79% for *P. ruminicola*, and 50% for *M. gottschalkii*). The rest of the unbounded reactions from the nucleotide degradation pathways were not fixed since they do not critically affect the biological significance of the models and are usually common in existing metabolic models (Fritzemeier et al., 2017). The initial manual curation process reconnected a number of blocked reactions in the models, which will be addressed in the subsequent sections. The reconciliation of cofactor inconsistency in *R. flavefaciens* model was performed by removing spurious duplicate reactions that caused unlimited ATP generation, thereby ensuring that all models are energetically consistent. The summary of model statistics before and after the manual curation are shown in Table 1. Supplementary Materials 1-3 contain the details of the microorganism models.

**Table 1:** Statistics for the draft and curated models of *R. flavefaciens, P. ruminicola*, and *M. gottschalkii*.

| Models | *R. flavefaciens* |  | *P. ruminicola* |  | *M. gottschalkii* |  |
| --- | --- | --- | --- | --- | --- | --- |
|  | Draft | Curated | Draft | Curated | Draft | Curated |
| Genes | 461 | 467 | 538 | 546 | 316 | 319 |
| Reactions | 982 | 1020 | 1041 | 1093 | 840 | 852 |
| Metabolites | 1025 | 1034 | 1067 | 1070 | 896 | 901 |
| Imbalanced Reactions | 251 | 0 | 49 | 0 | 36 | 0 |
| Unbounded Reactions | 38 | 8 | 54 | 12 | 16 | 8 |
| Blocked Reactions | 509 | 460 | 484 | 468 | 342 | 327 |
| Cofactor inconsistency | yes | no | no | no | no | no |

### Metabolic interactions in the microbial community

Individual metabolic models of the three species were integrated into an interacting community framework. This *in silico* community model was simulated at the growth condition estimated for a standard-sized domestic cow (see details in methods section). The simulated community biomass flux (0.513 h^−1^) is comparable to the experimentally observed values of passage/dilution rates in the rumen (Goetsch and Galyean, 1982;Tellier et al., 2004), which varied between 0.043 h^−1^ to 1.0 h^−1^ depending on dietary regimen and other factors. Dilution rate or passage rate of ruminal content is defined as the rate at which the digesta leaves a compartment in the gut (in this case, rumen) (Goetsch and Galyean, 1982;Tellier et al., 2004). It is important that the rumen microbiome growth rate (the community biomass flux) is within the margin of experimentally observed dilution rate because a stable community needs to survive and grow at a reasonable rate to perform its necessary role in host nutrition and pathophysiology despite constant washout events like fecal secretion. The extent and directionalities of the major metabolic exchanges in the community is shown in Figure 1. Complex plant carbohydrates and proteins are utilized by *R. flavefaciens* and *P. ruminicola*, and the short-chain fatty acids (SCFAs) are absorbed by the rumen epithelium. *M. gottschalkii* accepts hydrogen, formate, carbon di-oxide from the other two members, and produces methane, which is released to the atmosphere. It should be noted that all the metabolic transaction may not be active in all physiological condition of the community. When optimizing for the overall community biomass, the flux distribution (such as the flux space for shared metabolites shown in Figure 1) is shaped in a way that optimizes the production of biomass and not for other carbon molecules like short-chain fatty acids or small sugars. At the same time, when optimizing for other community objectives like production of SCFAs, some of the inactive fluxes including these metabolic transactions can become active. A comparative analysis of the effect of different community objective functions is presented in subsequent sections.

**Figure 1:**
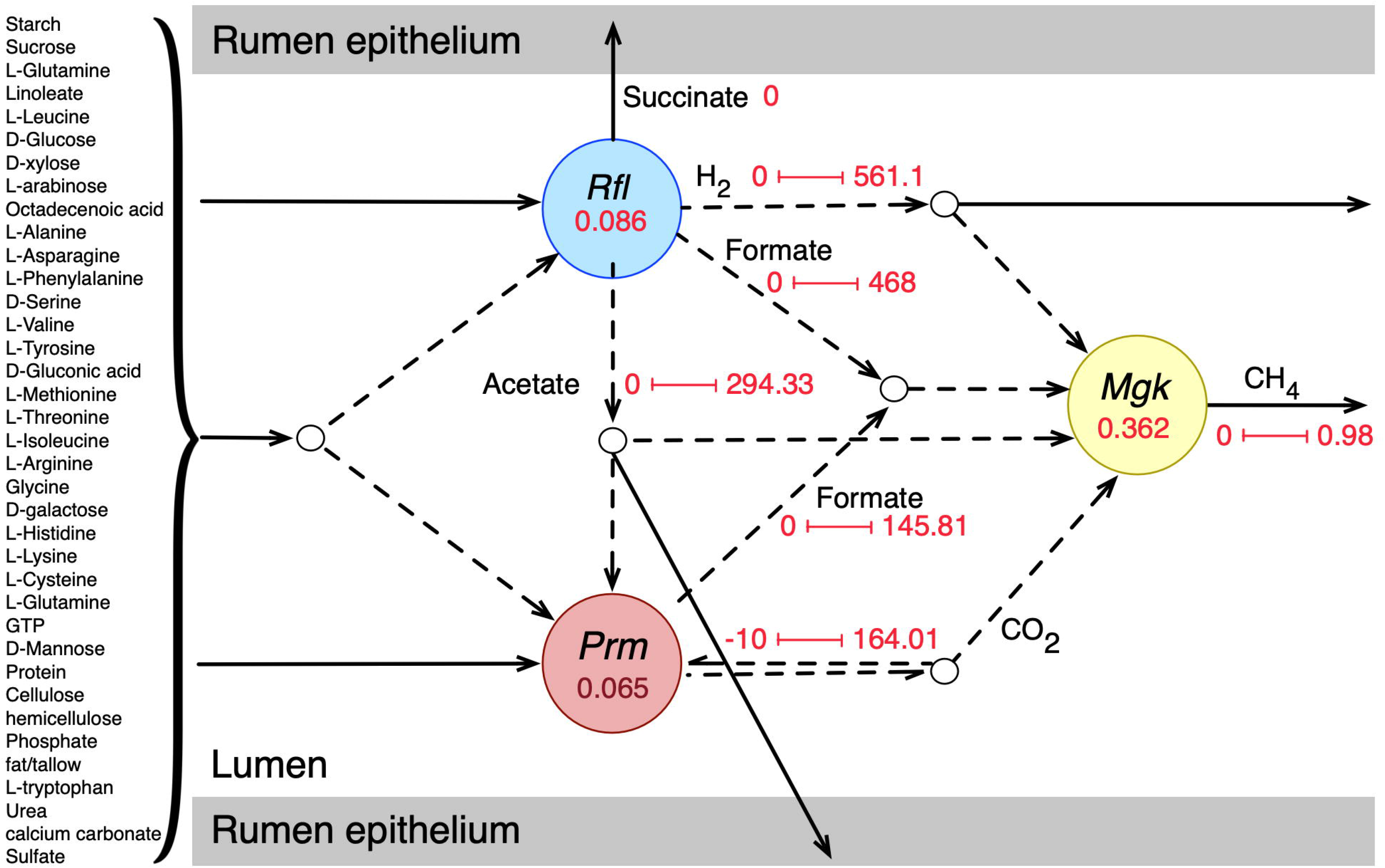
Initial community simulation results showing the interactions between the bacterial and archaeal members. **Rfl, Prm**, and Mgk represent *R. flavefaciens, P. ruminicola* and *M. gottschalkii*, respectively. The numbers inside the circles for each microbe represent the biomass flux (growth rate) of the respective microbe (hr^−1^). The arrows represent metabolic fluxes in mmol/gDCW.hr (dashes for inter-species/shared metabolites and solids for transfer to and from the rumen epithelium). The numbers along the arrows represent the minimum and maximum flux values.

### *De novo* interactions

The broad range of metabolic capabilities of different microorganisms in the rumen ecosystem is indicative of numerous inter-species interactions in both metabolic, signaling, and regulatory level. While the major functions of Bacteroides, Firmicutes and Archaea, and the interplay between them is commonly known (Hungate, 1975;Wolin, 1979;Flint, 1997;Raizada et al., 2003;Henderson et al., 2015;Nagaraja, 2016;Anderson et al., 2017;Liu et al., 2017), the current knowledge is only partial. As described in the methods section, a detailed step-by-step procedure was employed that identified 22 potential metabolic transactions. The identified *de novo* interactions are shown in Figure 2. In addition to the transfer of sugar monomers from *R. flavefaciens* to *P. ruminicola*, a number of co-enzymes and vitamins were found to be exchanged between *M. gottschalkii* and other members. Model statistics after filling the gaps using the GapFind-GapFill algorithms are shown in Table 2.

**Figure 2:**
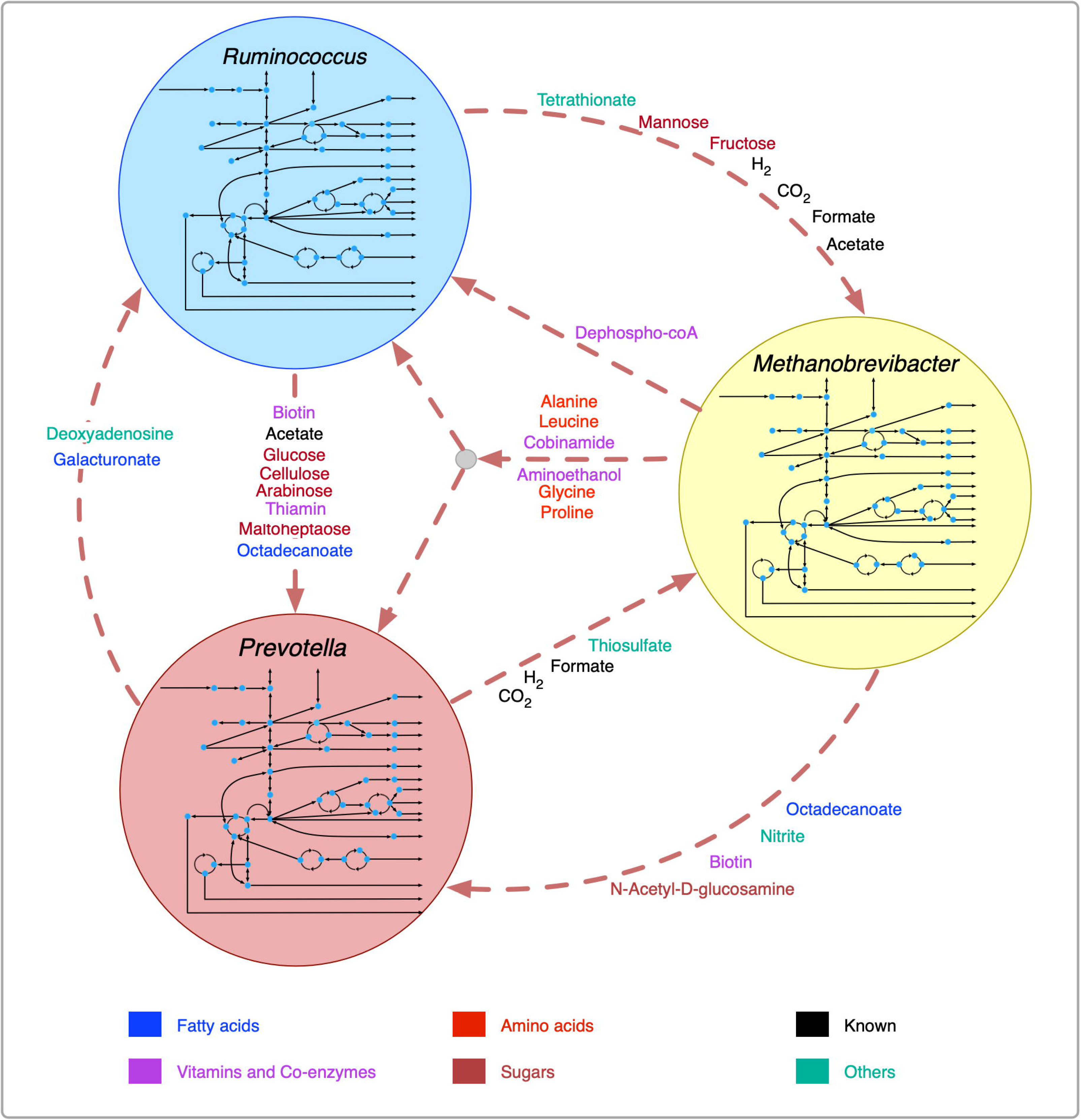
Identified *de novo* interactions in the community. Metabolites in black text were previously known to be exchanged, metabolites in color text are identified in this work. A cartoon inside the circles shows the main pathway map for each organism.

**Table 2:** Model statistics after filling the gaps using GapFind-GapFill.

| Models | *R. flavefaciens* | *P. ruminicola* | *M. gottschalkii* |
| --- | --- | --- | --- |
| Genes | 517 | 570 | 341 |
| Reactions | 1058 | 1143 | 909 |
| Metabolites | 1047 | 1110 | 924 |
| Imbalanced Reactions | 0 | 0 | 0 |
| Unbounded Reactions | 8 | 12 | 8 |
| Blocked Reactions | 396 | 389 | 242 |

The GapFill optimization procedure was repeated after the metabolic functions from the viruses were incorporated into the metabolic models of each of the microbes. Among the 22 identified de novo interactions as mentioned earlier, 20 were retained upon gap filling after adding these virome functionalities. The transfer of D-mannose and fructose from *R. flavefaciens* to *M. gottschalkii* did not appear in these gap filling results.

### Viral auxiliary metabolic genes and shifts in flux distributions in the metabolic models

Upon filtering of more than 3000 candidate proteins from BLAST search results for high bit score and expectancy values (<10^−34^), the models of *R. flavefaciens, P. ruminicola*, and *M. gottschalkii* were amended with 29, 26, and 18 metabolic reactions, respectively. For a detailed result of the alignment search see Supplementary Material 4. The reactions included additional metabolic functions and also novel metabolic capacities in the existing models. The addition of viral reactions resulted in significant changes in flux distributions. Up to 11% of reaction in all three of the models (See Supplementary Material 5 for details) had increased their flux space significantly, either by decreasing the minimum flux or by increasing the maximum flux or both. The reaction fluxes that had changed their range by at least one standard deviation were considered at this stage. Overall, all the reactions that had a changed flux distribution were relaxed in AMG-amended models compared to the post-gapfilled models. Figure 3 illustrates the change of flux space upon addition of viral AMGs.

**Figure 3:**
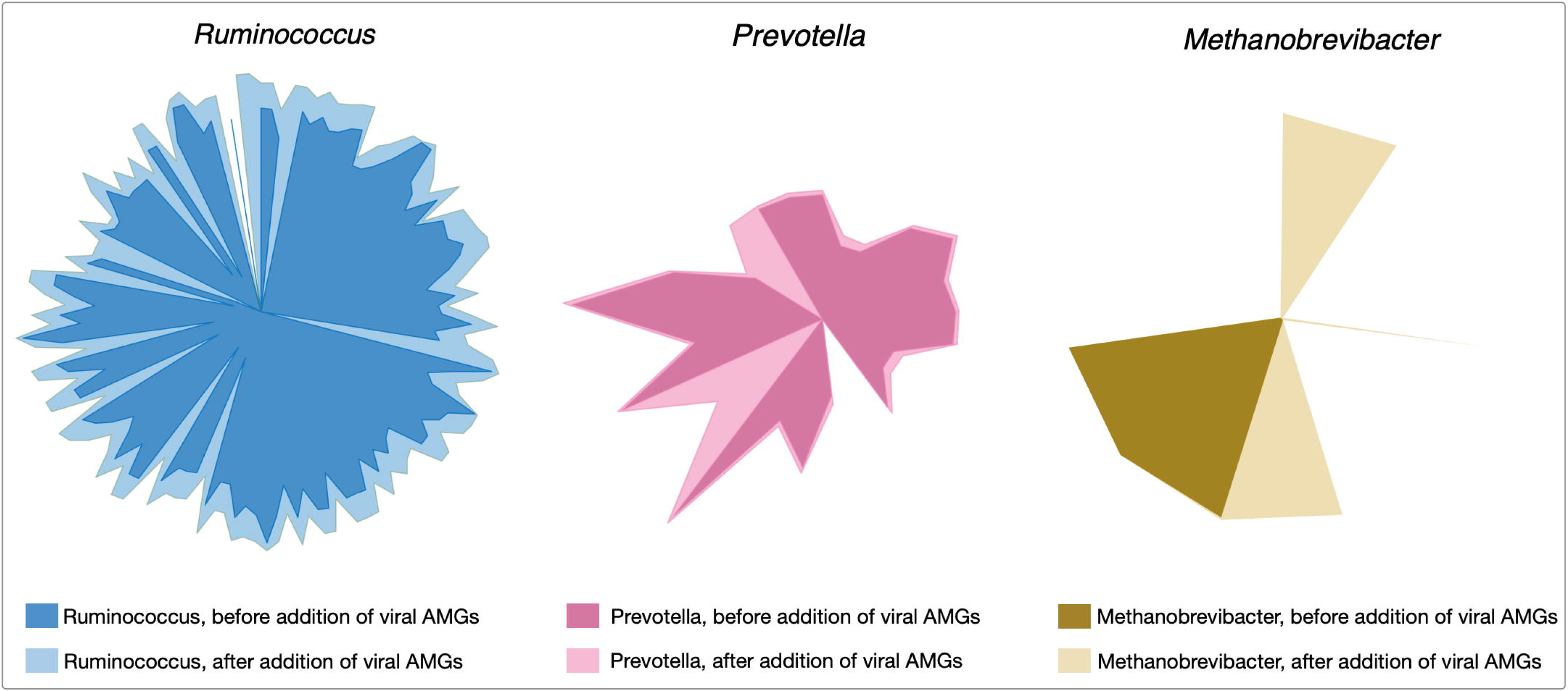
Changes in flux space after viral AMGs were added to the metabolic models. The lighter shade for each of the colors represent the relaxed flux space after addition of AMGs in each of the microbial metabolic models.

### Variability of the metabolic fluxes under different community objective functions

The metabolic spaces of the community members were assessed under different community objective functions after viral AMGs were added to the host models. To simulate different functions of the rumen microbial community, *i.e*., i) growth, ii) SCFA production, iii) feed utilization, iv) methane and carbon-di-oxide release, and v) small sugar molecule production, the appropriate community-level objective functions were chosen and optimized for (see Figure 4A for details). Statistical analysis was performed to investigate the deviation of flux ranges under these five objective functions. Approximately 44%, 18%, and 6% of the reactions from *R. flavefaciens, P. ruminicola*, and *M. gottschalkii* models, respectively, had non-zero standard deviation among their flux ranges in these conditions (See Supplementary Material 5 for details). However, under these five objective functions, the variation of flux distributions across the three community members was very small (with average standard deviation of 0.0001 and the range between 0 and 0.68). As representatives of the corresponding flux spaces, Figure 4B, 4C, and 4D show the distribution/density functions of the standard deviations of the exchange fluxes (i.e., both uptakes and imports) from all of the three community members. From this analysis it is evident that while viral AMGs play an important role in relaxing the flux ranges, the overall flux space of the intracellular and extracellular reactions in a community are very robust against the choice of a community level objective function.

**Figure 4:**
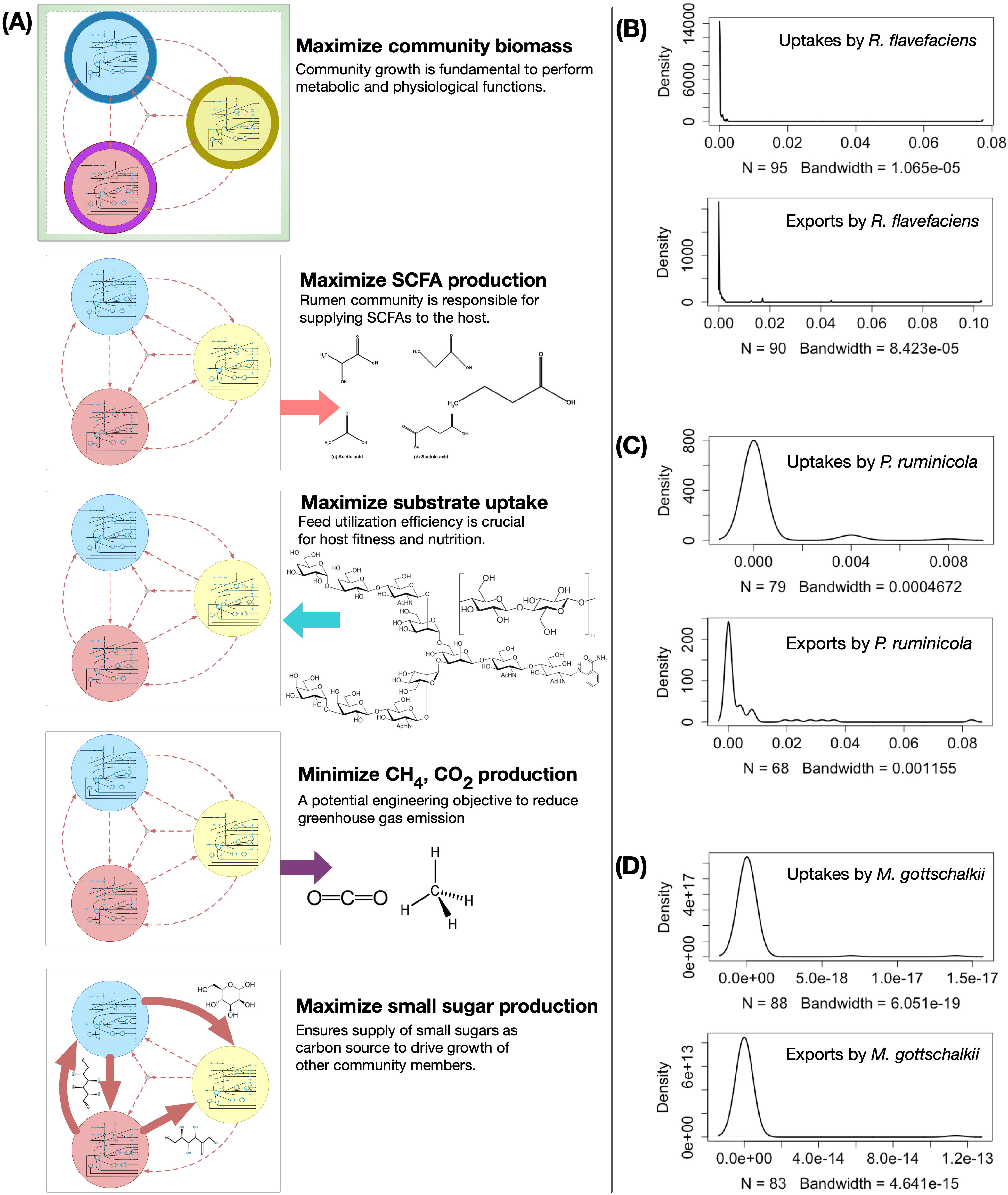
Variability in metabolic fluxes under different community objective functions. (A) Visual representation showing the choice of different community-level objective functions. The density functions (right) show the insignificance of the variations in exchange flux space upon optimizing for different objective functions, *i.e*, maximizing total community biomass, maximizing total Short-chain Fatty Acids (SCFA) production, maximizing total complex carbohydrate uptake, minimizing total Methane and carbon-di-oxide production, and maximizing total sugar production by the community: (B) *R. flavefaciens*, (C) *P. ruminicola*, and (D) *M. gottschalkii*. The bandwidth is the standard deviation of the smoothing kernel of the density function. N is the number of uptake and export fluxes for each of the organisms.

## Discussion

### Individual model performance and community interactions

The reconstructed and curated genome-scale metabolic models of *P. ruminicola*, R. *flavefaciens*, and *M. gottschalkii* were evaluated for fitness and species-specific metabolic capacities. Each of the metabolic models were simulated for growth and energy production at the growth condition estimated for a standard-sized domestic cow (see methods). The simulated growth rates were 0.086 hr^−1^, 0.065 hr^−1^ and 0.362 hr^−1^ for *R. flavefaciens, P. ruminicola*, and *M. gottschalkii*, respectively. While an experimental growth rate observation for these microbes are not available, the growth rates are comparable to the reported dilution rates in the rumen (Huntington et al., 1981; Estell and Galyean, 1985). The degradation of plant cellulose, biosynthesis of branched chain amino acid and short-chain fatty acid (SCFA) and production of hydrogen were predicted by *R. flavefaciens* model and were in agreement with experimental observations (Helaszek and White, 1991; Flint et al., 2008;Zheng et al., 2014). Protein digestion by *P. ruminicola* and methane production from carbon di-oxide and hydrogen consumption by *M. gottschalkii* were also validated (Wallace et al., 1997;Qiao et al., 2014;Henderson et al., 2015;Seedorf et al., 2015).

The representative community successfully captured most of the known metabolic interactions with the three functional guilds of microbes, as is evident from the flux values of the shared metabolites (Figure 1). While each of the community members were tested for known metabolic contribution in the community, a community simulation with total biomass as the biological objective rendered some of the metabolic transactions inactive. For example, even though the production of succinate and propionate by *R. flavefaciens* and *P. ruminicola* were validated, the maximization of community biomass did not drive the production of those SCFAs for growth. It should be noted that despite both *P. ruminicola* and *M. gottschalkii* being known and tested *in silico* for carbon di-oxide production in rumen, the community simulated showed that only *P. ruminicola* produced all the carbon di-oxide that was released from the rumen. This shows that metabolic exchanges can have separate dynamics based on community fitness criteria set during the optimization process.

### Insight into *de novo* community interactions

The predicted interactions between *R. flavefaciens* and *P. ruminicola* include the transfer of various small sugar monomers and fatty acids (as shown in Figure 2). These interactions are highly warranted given the metabolic functions each of these organisms perform (Wallace, 1992;Nagaraja, 2016). Both of these cellulolytic organisms contribute towards breakdown of complex plant material and the production of SCFAs and small sugar molecules for the host and other members in the community. Several NMR and GC-MS based metabolomic analyses of the ruminal fluid show the secretion of glucose, mannose, acetate, formate, and maltoheptaose in the rumen (Saleem et al., 2012;Saleem et al., 2013), which match our observations. These interactions build mutualistic and commensal relationships in the rumen ecosystem. *Prevotella* is also known for its proteolytic functions and high dipeptidyl peptidase activity (Nagaraja, 2016), which is demonstrated by the consumption and degradation of amino acids in the community simulation. This observation agrees to the meta-transcriptomic analysis by Li and Guan (Li and Guan, 2017), where they found downregulated amino acid synthesis and upregulated alanine, aspartate and glutamate catabolism in *Prevotella*, and the metabolomic study by Wang *et al*. (Wang et al., 2019), where a co-occurrence network analysis among the microbiota and metabolites showed positive correlation between *Prevotella* and several amino acids. Li and Guan (Li and Guan, 2017) also showed downregulated fatty acids synthesis pathway in *Prevotella*, which can be complemented by the uptake of fatty acids like octadecanoate produced by *R. flavefaciens* and *M. gottschalkii. Prevotella* also lacks the capability of glycan degradation (Li and Guan, 2017), which provides the explanation of our *in silico* observation why glycans are degraded by *R. flavefaciens* and then the degradation products are taken up by *Prevotella. Methanobrevibacter* acts as a sulfate reducer and a hydrogenotroph (Samuel and Gordon, 2006;Hansen et al., 2011;Zheng et al., 2014;Nagaraja, 2016), which can be attributed to its consumption of thiosulfate, formate, and hydrogen. The consumption of hydrogen by *Methanobrevibacter* may potentially facilitate in the extraction of energy from nutrients and increasing digestion efficiency by redirecting the ruminal fermentation towards more oxidized end products (Armougom et al., 2009). Previous studies have inferred an absence of Glycosaminoglycan degradation functions in *M. gottschalkii* (Li and Guan, 2017), which explains our observation of acetyl glucosamine secretion by *M. gottschalkii* and subsequent consumption by *P. ruminicola*. At the same time, the role of *Methanobrevibacter* in supplying a number of important amino acids, vitamins and co-enzymes to the other organisms was observed, which was not reported before the current study. These roles suggest that *Methanobrevibacter* is a major player in the rumen ecosystem and warrants further studies on its role in specificity and efficiency of bacterial digestion in rumen.

### Relaxation of metabolic bottlenecks in presence of viral auxiliary metabolic genes

The addition of Auxiliary Metabolic Genes (AMGs) from the viruses associated with each of the host community members had profound impact on shifting and relaxing the flux spaces in the host organisms. As seen in Fig. 3, in *R. flavefaciens* 127 reactions were observed to have wider flux ranges after virome related AMGs were added, compared to the gap-filled model. The pathways that were relaxed due to viral metabolic genes are Calvin-Benson cycle (CBB), amino acid biosynthesis, Sugar utilization, nucleoside catabolism, fermentation, glycolysis/gluconeogenesis, cofactor biosynthesis, single carbon metabolism (tetrahydropterines), and various transport mechanisms. In addition to that, Pentose Phosphate Pathway (PPP) carried higher flux compared to the gapfilled model. Therefore, viral AMGs not only add some additional metabolic functions to host but also complement and relax some key pathways that drive the fitness of the host. In *P. ruminicola*, 25 reactions were relaxed after virome related AMGs were added, compared to the gap-filled model. The pathways that were relaxed due to viral metabolic genes are folate biosynthesis, energy production, cofactor synthesis, and glycogen synthesis, which are important for boosting the energy production in the microbes and the generation of reducing power. It has also been previously hypothesized that these phenomena aid in viral replication (Anderson et al., 2017). In *M. gottschalkii*, 11 reactions had wider flux ranges after viral AMGs were added. The relaxed pathways include amino acid biosynthesis and degradation, and nucleotide metabolism. In addition, it was observed that coenzyme biosynthesis, glycine and serine degradation, diaminopimelic acid (DAP) pathway for lysine synthesis, methanogenesis, methionine biosynthesis, peptidoglycan biosynthesis, pentose phosphate pathway, and purine and pyrimidine conversion pathways carried higher fluxes compared to the post-gapfilled model. These show the important role of viral AMGs in cell growth and replication for the host (*M. gottschalkii*).

The inclusion of viral AMGs as metabolic functions resulted in noticeable changes in the inter-species transports, as manifested in the change in the flux ranges of the shared metabolite transactions (See Supplementary Material 5 for details). The overall changes in metabolic interactions among *R. flavefaciens, P. ruminicola*, and *M. gottschalkii* are shown in Figure 5. The transfer of mannose and fructose sugars to *M. gottschalkii* from *R. flavefaciens* was not identified as interactions, which ultimately resulted in reduction of the number of *de novo* interactions in the community to 20 when viral AMGs were present. Upon inspection of the functionalities coming from the phages associated with *M. gottschalkii*, it became apparent that the D-glyceraldehyde-3-phosphate glycolaldehyde transferase and D-mannose-6-phosphate aldose-ketose-isomerase system that drives the conversion between fructose and mannose became unnecessary in the presence of virome functionalities in the models. At the same time, the phosphofructokinase/sedoheptulokinase (EC 2.7.1.14) enzymes, including Sedoheptulose 1,7-bisphosphate D-glyceraldehyde-3-phosphate-lyase and ATP:Sedoheptulose 7-phosphate 1-phosphotransferase could shift the carbon flux from glycolysis (via fructose-6-phosphate) to Pentose Phosphate pathway through tetrose sugars and therefore facilitate the adaptation of *M. gottschalkii* even without the support of sugars from *R. flavefaciens*. The ability to maintain species fitness without these interactions leads to better community fitness in situations when these interactions are threatened (e.g., antibiotic treatment).

**Figure 5:**
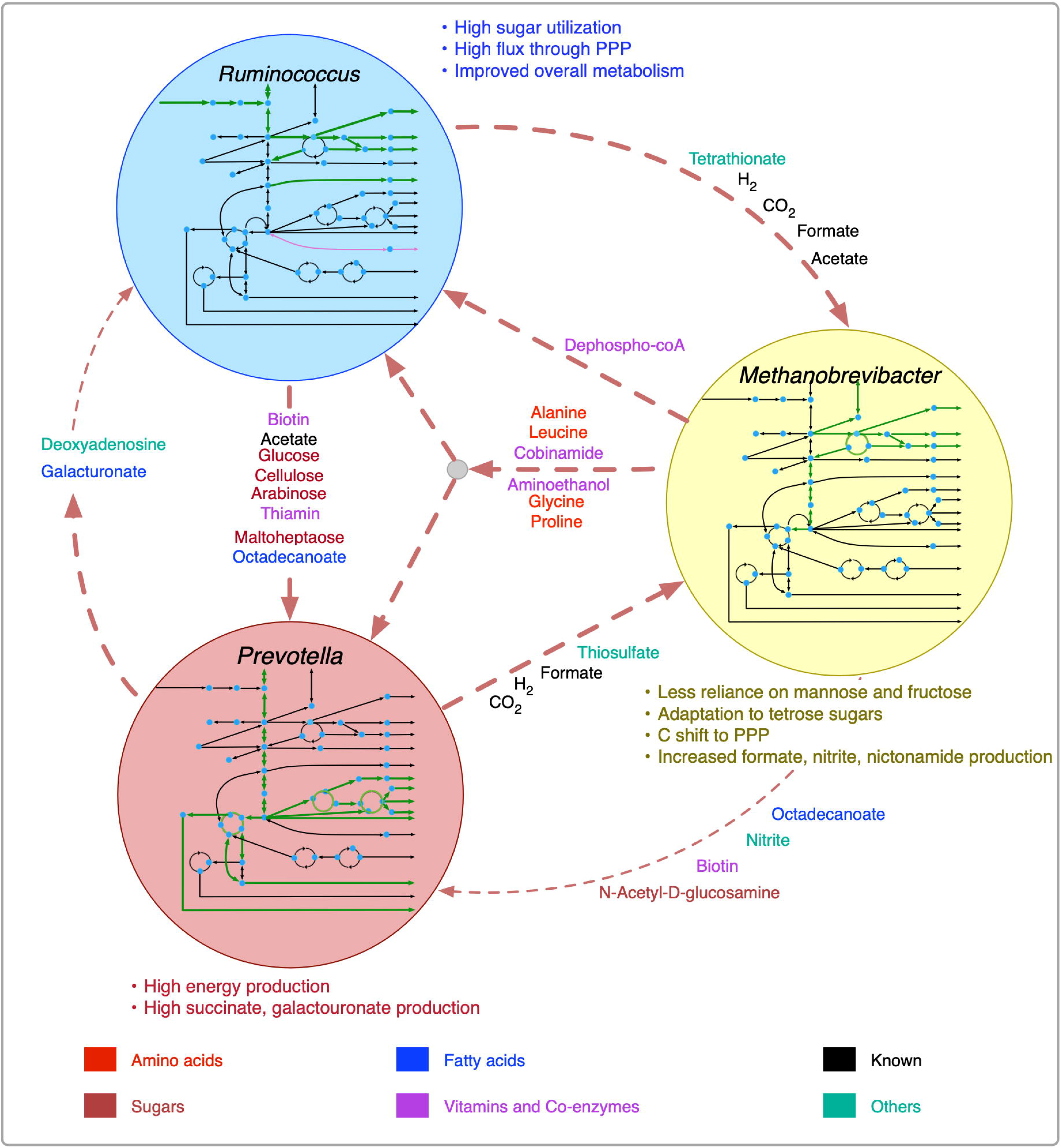
Shifts in metabolism and inter-species interactions after the inclusion of viral auxiliary metabolic genes. Inside the circles for each organism, increase in pathway fluxes are shown in thicker green lines and decrease in pathway fluxes are shown in purple lines. Decreased metabolic transactions are shown in thinner dashed lines.

### Robust community behavior under different community objectives

Although living organisms have evolved to maximize their survival in varying environments, it is difficult to assess the primary driver of the flux distributions of primary metabolism (Burgard and Maranas, 2003a). Disagreement is prevalent among the scientific community as to whether metabolic flux in a network is distributed to satisfy optimal biomass production, maximum energy generation, or most efficient utilization of substrates. It is especially true for a naturally occurring microbial community such as the rumen ecosystem in which thousands of microbial, viral, fungal, and archaeal species co-evolved with different metabolic functions and interplay among one another. Therefore, maximizing community biomass may not always be the best choice for an objective function. However, choosing the best community objective is not as straightforward as it seems. With that in mind, the variation of flux ranges under different community objective functions was studied. From Figure 4 (also Supplementary Material 5), it is observed that the choice of a community objective has negligible effect while simulating the possible metabolic capacities (flux space) of the community members. While there have been attempts at optimizing a number of different objective functions for prokaryotic microorganisms, optimizing for growth often seems realistic for both eukaryotic and prokaryotic organisms (Burgard and Maranas, 2003a). Due to the lack of knowledge about the overall goal of the rumen community, maximizing community biomass is a logical choice, given that a stable community in the rumen needs to survive and grow at a reasonable rate to perform its necessary role in host nutrition and pathophysiology despite constant washout events like fecal secretion.

### Moving ahead: challenges and future prospects

In summary, we developed a workflow to elucidate novel interactions between participating members of a complex community and its phages. Our *in silico* predictions of metabolite exchange between three members of rumen microbiome agrees with the current knowledgebase about their metabolic functionalities and roles in the rumen ecosystem and also with recently published multi-omics datasets. We also identified possible viral auxiliary metabolic genes associated with the three members in our community that reinforced their metabolic capacities and help in relaxing several bottlenecks in the metabolic network models. This was manifested by the enhancement of reaction fluxes in important metabolic pathways in the models and the metabolic robustness achieved by the microbes. Our community metabolic model serves to discover unidentified metabolite transactions and answer key ecological questions of ruminant nutrition through virome-microbiome interactions, while promising to address important biological aspects of ruminant nutrition and greenhouse emission. The development of additional bioinformatics tools, advancement in high-throughput sequencing technologies and cultivation-independent ‘omics’ approaches may drive further development of new mathematical frameworks for analyzing rumen ecosystems. *In vitro* cell culture systems and *in situ* experimentation of simplified microbial, viral, and fungal communities in gnotobiotic bovine rumen to capture spatiotemporal community dynamics will enhance our analyzing power to a greater extent. Therefore, what we need is a combination of computational and experimental efforts to enrich the current knowledgebase regarding in situ metabolic and taxonomic profiling, species identification and characterization, annotation, and advanced tools to accommodate for large-scale data analysis and integration, which will bring success in deciphering the complexity of this ecosystem.

## Methods

### Model reconstructions and refinements

The initial draft genome-scale metabolic reconstructions of *P. ruminicola, R. flavefaciens*, and *M. gottschalkii* were created and downloaded from the Modelseed biochemical database (in September 2017). The models included reactions for central carbon metabolism, secondary biosynthesis pathway, energy and cofactor metabolism, lipid synthesis, elongation and degradation, nucleotide metabolism, amino acid biosynthesis and degradation. Flux Balance Analysis (FBA) was employed during model testing, validation, and analyzing flux distributions at different stages of our work (Varma and Palsson, 1993;Varma and Palsson, 1994;Oberhardt et al., 2009a). The optimization formulation, in its most common form, is given below.

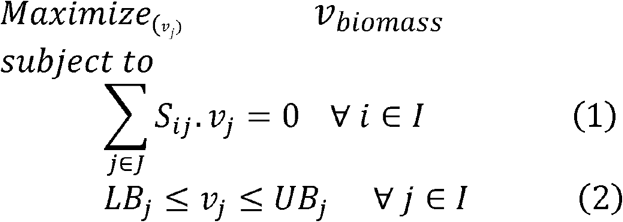

Here, *I* and *J* are the sets of metabolites and reactions in the metabolic model, respectively. *S_ij_* is the stoichiometric coefficient of metabolite *i* in reaction *j* and *v_j_* is the flux value of reaction *j*. Parameters *LB_j_* and *UB_j_* denote the minimum and maximum allowable fluxes for reaction *j*, respectively. *v_biomass_* is the flux of the biomass reaction which mimics the cellular growth yield.

### Curation of metabolic models

#### Correcting reaction imbalances

For balancing the reactions imbalanced in protons, we checked for the protonation state consistent with the reaction set in the draft model and performed addition/deletion of one or multiple protons on either the reactant or the product side. For the remaining imbalanced reactions, we corrected the reaction stoichiometry in order to ensure that the atoms on both sides of the reactions balance out.

#### Identifying and eliminating thermodynamically Infeasible Cycles

One of the limitations of constraint-based genome-scale models is that the mass balance constraints only describe the net accumulation or consumption of metabolites, without restricting the individual reaction fluxes. While biochemical conversion cycles like TCA cycle or urea cycle are ubiquitous in a metabolic network model, there can be cycles which do not consume or produce any metabolite. Therefore, the overall thermodynamic driving force of these cycles are zero, implying that no net flux can flow around this cycle (Schellenberger et al., 2011). To identify Thermodynamically Infeasible Cycles in our model, we turned off all the nutrient uptakes to the cell and used an optimization formulation called Flux Variability Analysis (FVA) which maximizes and minimizes each of the reaction fluxes subject to mass balance constraints (Mahadevan and Schilling, 2003). The reaction fluxes which hit either the lower bound or upper bound are defined as unbounded reactions and were grouped together as a linear combination of the null basis of their stoichiometric matrix. To eliminate the cycles, we either removed duplicate reactions, turned off lumped reaction or selectively turned reactions on/off based on available cofactor specificity information (see Supplementary Material 6 for details). The mathematical formulation of Flux Variability Analysis (FVA) is given below. *v_app-obj, threshold_*(from constraint 3, which is optional) is a predetermined threshold value of the appropriate objective flux *v_app-obj_* to ensure that the feasible flux space satisfy the targeted value.

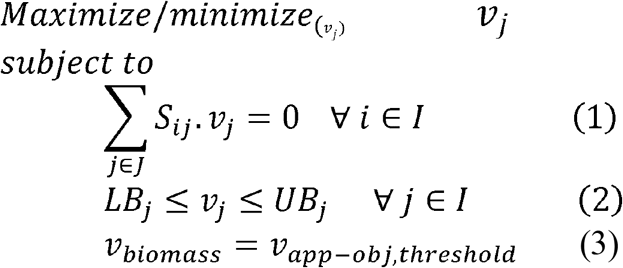

#### Ensuring known metabolic functions

Each of the three metabolic models were checked for the capacity to produce biomass and metabolites they were known to produce (Helaszek and White, 1991;Wallace et al., 1997;Flint et al., 2008;Qiao et al., 2014;Zheng et al., 2014;Henderson et al., 2015;Seedorf et al., 2015). To ensure these metabolic functionalities, the reactions missing in metabolic pathways were systematically identified and added manually from biochemical databases (Kanehisa and Goto, 2000;Apweiler et al., 2004;Henry et al., 2010) after an extensive search for each of the missing enzyme activities in related organisms such as *Ruminococcus albus, Bacteroides thetaiotaomicron*, and *Methanobrevibacter smithii* for *R. flavefaciens, P. ruminicola* and *M. gottschalkii*, respectively. A missing metabolic function was only added if the genes between any pair of organisms were found to be orthologous and amending the models did not result in an increase in the number of thermodynamically infeasible cycles.

### Community formation and simulation

Once the individual microbe models were curated, they were integrated to form a community model using existing optimization framework, namely OptCom (Zomorrodi and Maranas, 2012). In this bi-level multi-objective optimization framework, the individual flux balance analysis problems for each community member are treated as inner-level optimization problem, and the community objective is optimized for in the outer-level problem. The mathematical description of the OptCom procedure is given below.

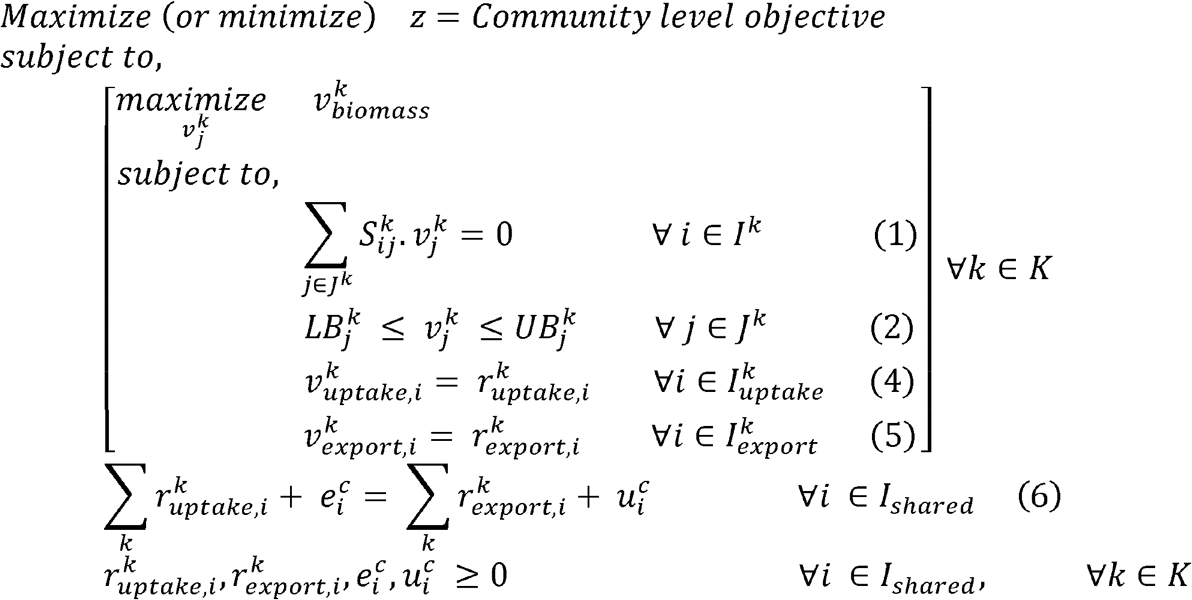

The inner problem(s) represents the steady-state FBA problem for each of the community members *k* and limits on uptake or export flux of a shared metabolite using the parameters 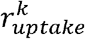 and 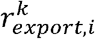, respectively, which are imposed by the outer problem. Constraint (6) in the outer problem describes a mass balance for each shared metabolite in the extra-cellular environment of shared metabolite pool, where the terms 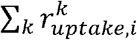 and 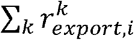 represent the total uptake and export of the shared metabolite *i* by community members, respectively. These constraints are the key equations for modeling the known metabolic interactions among participants of the community. When further information on *de novo* interaction became available, they were also incorporated using these outer-level constraints. Incorporating the known interaction in the community, a rumen ecosystem for a 1000 lb cow using the three-member simplified community was simulated. The community nutrient uptakes were designed based on the data obtained from previous experiments (Anderson et al., 2017) and details are given in Supplementary Material 7.

### Identification of viral auxiliary metabolic genes

In addition to the syntrophic, mutualistic, and competitive microbial interactions, viruses impact microbial populations through cell lysis and reprogramming of host metabolism by Auxiliary Metabolic genes (AMGs). Most of the bacterial species in the cattle rumen has its own associated viruses (Berg Miller et al., 2012;Ross et al., 2013;Parmar et al., 2016). To identify the viral AMGs, the viruses/phages associated with each of the community members were searched for. Miller *et al* (Berg Miller et al., 2012) suggested 13 different phages associated with *R. flavefaciens, P. ruminicola* and *M. gottschalkii*. A local alignment search (BLAST) of the viral proteomes downloaded from several databases (Apweiler et al., 2004;UniProt, 2007;Hubisz et al., 2011;Zhou et al., 2011;Brister et al., 2015;Arndt et al., 2016) to The National Center for Biotechnology Information (NCBI) non-redundant proteins sequence database (O’Leary et al., 2016) was performed. The search yielded more than 3000 candidate proteins, which were filtered for expectancy values (<10^−34^).

### Identification of unknown interactions and bridging of network gaps

One of the key limitations of any genome-scale reconstructions is the gaps in the resultant metabolic network models. Any such gaps from a specific model could be reconciled by borrowing the required metabolic functionalities from the models of neighboring organism. For each of the metabolic models in the rumen community, a set of Mixed-integer Linear Programming (MILP) optimization procedures (named GapFind and GapFill) was used to identify and eliminate network gaps in these reconstructions (Satish Kumar et al., 2007). To identify unknown inter-species interactions in the community, a protocol was developed that takes each of the suggestion from the GapFill algorithm and performs a series of tests (shown in Figure 6) to categorize the results as unacceptable, possible inter-species interactions, or acceptable solutions to bridge network gaps. The metabolites that cannot be produced or consumed in a network are called the problem metabolites. For the problem metabolites in each of the member species in the community, we performed three separate set of gap filling procedures. Two of them used the other community members as the source database, and one used the Modelseed biochemical database (Henry et al., 2010) as the source database (as of February 2018). For each of the gap filling solutions, it was tested if the solution was obtained from another member in the community or the Modelseed database. If it was from the Modelseed database, it was tested for possible transport mechanisms (Ren et al., 2007), likelihood based on presence in taxonomically related organisms (with prioritizing organisms in the same lower taxonomic levels), and whether the solution created thermodynamically infeasible cycles, and finally that solution was either accepted or rejected. Similarly, for a gap filling suggestion coming from another member in the community, the same sets of checks were performed, and the solution was either accepted or rejected. However, for these gap filling suggestions, if the solution was a transport mechanism and the target organism was known to have a transport mechanism as per the current knowledgebases (Ren et al., 2007), it was concluded to be a *de novo* identification of a potential interaction that exists in the community. It should be noted that for each transport reaction, strong bioinformatic evidence or clear orthologue of experimentally characterized transporter in a closely related organism was ensured before it was accepted as a gapfill solution. When the individual metabolic models for each of the microbes were augmented with viral AMGs, the interaction identification procedure (shown in Figure 6) was repeated to compare the shifts in inter-species interactions. Therefore, the gapfiling procedure and the protocol described above served two very important steps in the model curation process by bridging network gaps as well as identifying possible metabolic interactions between organisms.

**Figure 6:**
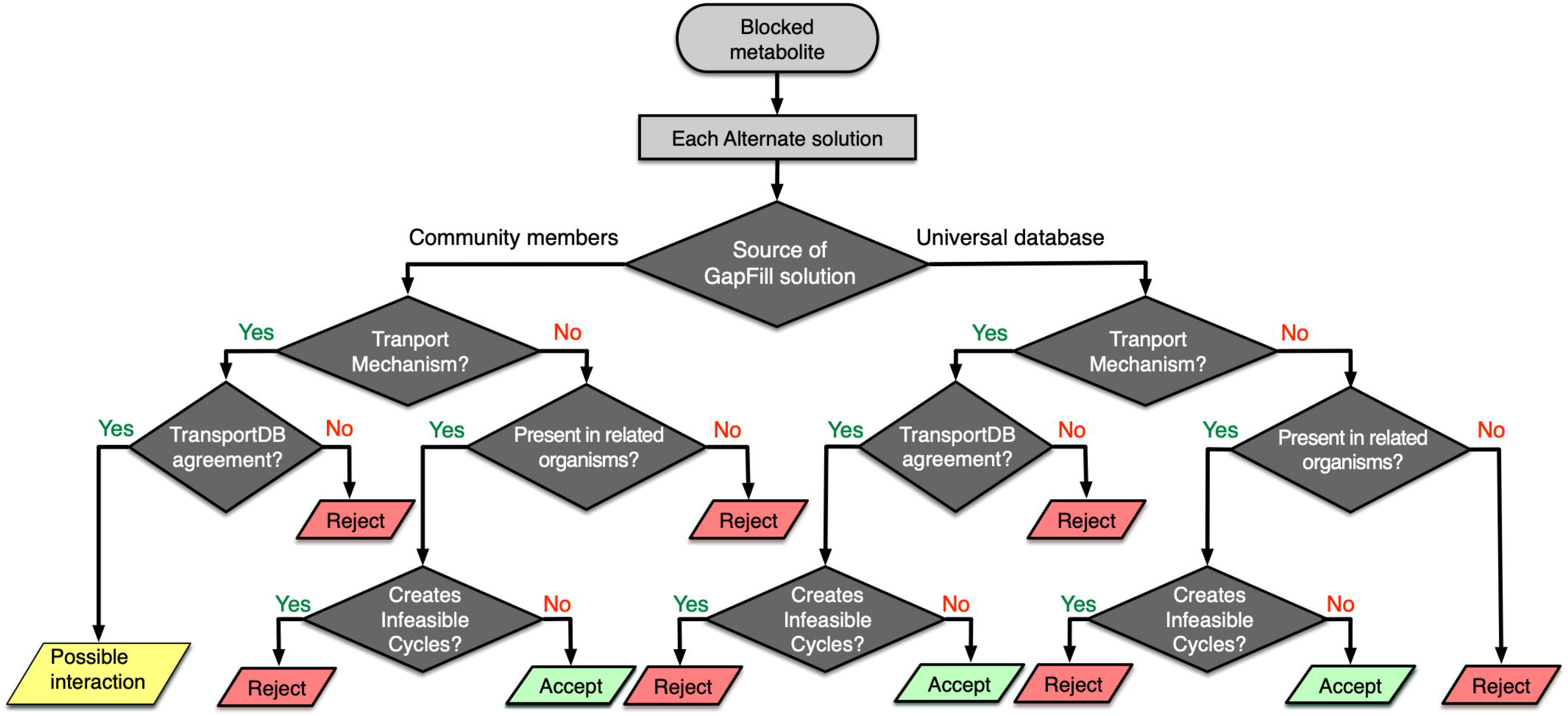
Workflow for identifying possible interspecies interactions and filtering GapFill suggestions.

## Acknowledgements

The authors would like to thank Mr. Wheaton Schroeder and Mr. Matthew Van Beek from Chemical and Biomolecular Engineering Department at the University of Nebraska-Lincoln for their help during the model curation step.

## Author contributions

RS and SF conceived the study. RS supervised the study. MI performed all the experiments and analyzed the results. MI, SF and RS wrote the manuscript. All authors read and approved the manuscript.

## Data Availability Statement

All datasets generated or analyzed for this study are included in the manuscript and the supplementary files.

## Supplementary Material

**Supplementary Material 1: Model files for *R. flavefaciens***

**Supplementary Material 2. Model files for *P. ruminicola***

**Supplementary Material 3. Model files for *M. gottschalkii***

**Supplementary Material 4. BLAST search results**

**Supplementary Material 5. Flux distributions in the models at different stages**

**Supplementary Material 6. Strategies for breaking infeasible cycles**

**Supplementary Material 7. Details of the community nutrient uptake calculations**

