## Supplementary Material 6 for "Metabolic Modeling Elucidates the Transactions in the Rumen Microbiome and the Shifts upon Virome Interactions"

**Strategies for fixing thermodynamically infeasible cycles**

To fix the thermodynamically infeasible cycles in the models, three distinct cases were addressed.

**Case 1: Duplicate reactions that run in opposite direction**

In this case the model contains duplicates of the same reaction, often one being irreversible and one being reversible. The cycle can be broken by removing or turning off one of the reactions, usually the irreversible one if no concrete thermodynamic information is available.

Example:

Phosphoglycerate dehydrogenase (PGCD):

nad_c[c] + 3pg_c[c] -> h_c[c] + nadh_c[c] + 3php_c[c]

and

Phosphoglycerate dehydrogenase reversible (PGCDr):

nad_c[c] + 3pg_c[c] <=> h_c[c] + nadh_c[c] + 3php_c[c]

Solution:

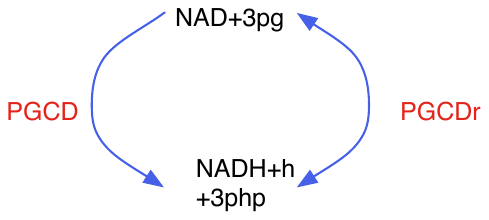

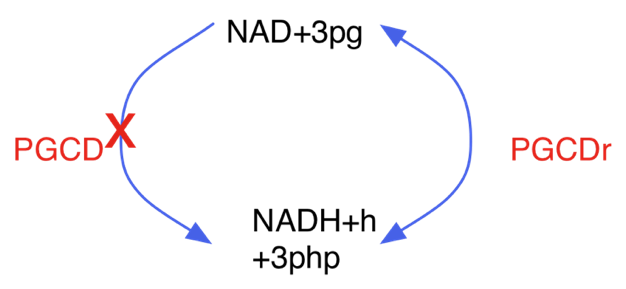
Turn off PGCD. Kept directionality of PGCDr as reversible.

**Supplementary Figure 1**. Example of fixing cycles involving duplicate reactions.

**Case II: Lumped reactions**

In this case, multiple reactions in a pathway are lumped together to represent the overall conversion. If both the individual reactions and the lumped reaction are present in the model, they can potentially create thermodynamically infeasible cycles. The cycle can be broken by removing or turning off the lumped reaction and assigning proper annotation information to the individual reactions.

Example:

Aconitase (ACONT): cit_c[c] -> icit_c[c]

Aconitase (half-reaction A, Citrate hydro-lyase, ACONTa): cit_c[c] <=> h2o_c[c] + acon-C_c[c]

Aconitase (half-reaction B, Isocitrate hydro-lyase, ACONTb): icit_c[c] <=> h2o_c[c] + acon-C_c[c]

Solution:

The lumped reaction (ACONT) can be turned off.

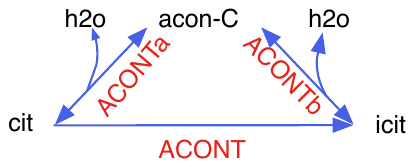

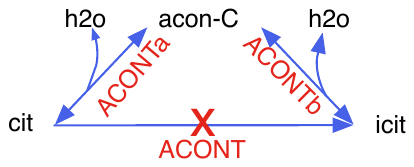

**Supplementary Figure 2**. Example of fixing cycles involving lumped reactions.

**Case III: Cofactor specificity**

In this case, the same biochemical conversion is carried out by different cofactors in the model, while in reality the organism only uses one of the cofactors. If the cofactor specificity information is available, the reaction with non-specific cofactor can be removed or turned off.

Example:

D-Ribitol-5-phosphate NAD 2-oxidoreductase (DR1ORx ):

nad_c[c] + dr5p[c] <=> h_c[c] + nadh_c[c] + ru5p-D_c[c]

and

D-Ribitol-5-phosphate NADP 2-oxidoreductase (DR1ORy):

nadp_c[c] + dr5p[c] <=> h_c[c] + nadph_c[c] + ru5p-D_c[c]

both catalyzes the conversion of D-Ribitol-5-phosphate to Ribulose-5-phosphate in S. aureus.

Solution:

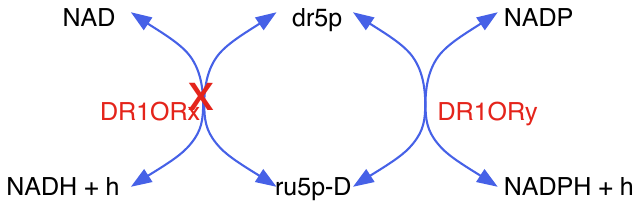
Upon extensive search for evidence in literature for cofactor specificity of S. aureus for this reaction, DR1ORx was turned off.

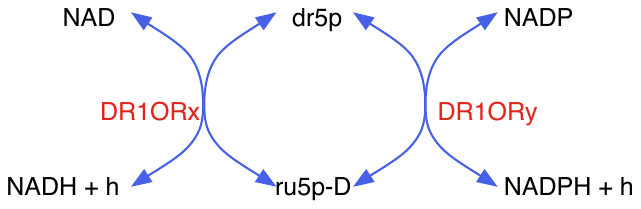

**Supplementary Figure 3**. Example of fixing cycles involving non-specific cofactors.
