## Supplementary Material 7 for "Metabolic Modeling Elucidates the Transactions in the Rumen Microbiome and the Shifts upon Virome Interactions"

**Details of the community nutrient uptake calculations**

**Converting the nutrient feed rates in mmol/h for adult cattle to the uptake flux for microorganisms in mmol/gDW.h.**

The average percentage of water in living cell is assumed to be 70%. Data about change in microbial organic matter in cattle rumen after feeding from Craig et al (Craig et al., 1987) was used to estimate the washout rate of microorganisms from the rumen (0.77 gDW/L.h). Dilution rate of the rumen is found to be varying between 1.4/day or (0.06 /h) and 0.32/h (Stokes et al., 1985). The usual volume of the rumen is ~70 L (Wolin, 1979). Therefore, the feed rates of the nutrients have been converted to nutrient uptake fluxes according to the following equation (starch uptake as an example). Table 2 lists the metabolites the calculated uptake rates.

$$v_{uptake, starch}= \frac{162.6 \frac{mmol}{h} \times0.32\frac{1}{h}}{0.77 \frac{gDW}{L.h} \times70 L}=\sim1 \frac{mmol}{gDW.h}$$

Starch is considered to be a dimer of glucose with a molecular weight ~348. Therefore, starch was fed to the models as two glucose molecules.

Table 2: Calculated nutrient uptake rates in the rumen community.

| Metabolites ID | Metabolite Name | % feed composition | uptake flux (mmol/gDWW.h) |
| --- | --- | --- | --- |
| cpd11657 | Starch | 12.530908 | 0.965423023 |
| cpd00076 | Sucrose | 11.43602571 | 0.834726132 |
| cpd00053 | L-Glutamine | 9.337021855 | 1.597816364 |
| cpd01122 | Linoleate | 6.815954262 | 0.610370885 |
| cpd00107 | L-Leucine | 6.021164448 | 1.148366562 |
| cpd00027 | D-Glucose | 5.723202176 | 0.794397609 |
| cpd01422 | D-xylose | 5.270935463 | 0.877945961 |
| cpd00224 | L-arabinose | 4.001040121 | 0.666427627 |
| cpd00035 | L-Alanine | 3.344362139 | 0.938846598 |
| cpd00132 | L-Asparagine | 2.115054647 | 0.400330802 |
| cpd00066 | L-Phenylalanine | 2.04736552 | 0.310015056 |
| cpd00550 | D-Serine | 1.90483548 | 0.317276246 |
| cpd00156 | L-Valine | 1.806927953 | 0.385856944 |
| cpd00069 | L-Tyrosine | 1.582322899 | 0.218417727 |
| cpd00060 | L-Methionine | 1.395084356 | 0.233929798 |
| cpd00161 | L-Threonine | 1.243982408 | 0.261179221 |
| cpd00322 | L-Isoleucine | 1.107191645 | 0.211165444 |
| cpd00051 | L-Arginine | 0.93620388 | 0.133660701 |
| cpd00033 | Glycine | 0.806283864 | 0.268595078 |
| cpd00348 | D-galactose | 0.782808808 | 0.075513967 |
| cpd00119 | L-Histidine | 0.772743767 | 0.124559013 |
| cpd00039 | L-Lysine | 0.578796694 | 0.098373937 |
| cpd00084 | L-Cysteine | 0.556558968 | 0.114920408 |
| cpd00053 | L-Glutamine | 0.470811116 | 0.080568485 |
| cpd00038 | GTP | 0.160980374 | 0.007734655 |
| cpd00138 | D-Mannose | 0.156561762 | 0.021731242 |
| cpd00052 | CTP | 0.140397548 | 0.00730785 |
| cpd00002 | ATP | 0.126879201 | 0.006289719 |
| cpd00062 | UTP | 0.114216846 | 0.005932755 |
| cpd00115 | dATP | 0.022409454 | 0.001147315 |
| cpd00357 | dTTP | 0.022000697 | 0.001147552 |
| cpd00241 | dGTP | 0.02042823 | 0.001012678 |
| cpd00356 | dCTP | 0.018873001 | 0.001016235 |
| cpd11746 | Cellulose | 2.082792938 | 0.320930655 |
| cpd29869 | hemicellulose | 3.319768937 | 0.002499522 |
| cpd00009 | Phosphate | 0.020151259 | 0.005244478 |
| cpd00065 | L-tryptophan | 0 | 0 |
| cpd00073 | Urea | 1.510934394 | 0.629166662 |
| cpd05097 | calcium carbonate | 1.461232604 | 0.598495465 |
| cpd00048 | Sulfate | 0.016419544 | 0.004273279 |

The growth medium metabolites in the models have been classified into three types: metabolites which are essential and limiting for growth, metabolites which are just limiting for growth, and metabolites that are not necessary at all or has no effect on growth. The uptake rates were converted to uptake fluxes in appropriate unit (mmol/gDCW.hr). The presence of metabolites in the model were compared to the list of metabolites obtained from the different diets. The following strategies were employed to design the feed for the individual and community metabolic models.

- For inorganic ions, the uptake values were set to big M (1000).

- If the nutrient is growth limiting and essential

- if the nutrient is found in the diet, then it’s uptake was set to the calculated values

- otherwise, uptake value of 1 mmol/gDW.h is used

- If the nutrient is not necessary at all

- if the nutrient is found in the diet, then it’s uptake was set to the calculated values

- otherwise, no uptake was allowed.
